# ∆^9^-tetrahydrocannabinol negatively regulates neurite outgrowth and Akt signaling in hiPSC-derived cortical neurons

**DOI:** 10.1101/440909

**Authors:** Carole Shum, Lucia Dutan, Emily Annuario, Katherine Warre-Cornish, Samuel E. Taylor, Ruth D. Taylor, Laura C. Andreae, Noel J. Buckley, Jack Price, Sagnik Bhattacharyya, Deepak P. Srivastava

## Abstract

Endocannabinoids regulate different aspects of neurodevelopment. *In utero* exposure to the exogenous psychoactive cannabinoid Δ^9^-tetrahydrocannabinol (Δ^9^-THC), has been linked with abnormal cortical development in animal models. However, much less is known about the actions of endocannabinoids in human neurons. Here we investigated the effect of the endogenous endocannabinoid 2-arachidonoyl glycerol (2AG) and Δ^9^-THC on the development of neuronal morphology and activation of signaling kinases, in cortical glutamatergic neurons derived from human induced pluripotent stem cells (hiPSCs). Our data indicate that the cannabinoid type 1 receptor (CB1R), but not the cannabinoid 2 receptor (CB2R), GPR55 or TRPV1 receptors, is expressed in young, immature hiPSC-derived cortical neurons. Consistent with previous reports, 2AG and Δ^9^-THC negatively regulated neurite outgrowth. Interestingly, acute exposure to both 2AG and Δ^9^-THC inhibited phosphorylation of serine/threonine kinase extracellular signal-regulated protein kinases (ERK1/2), whereas Δ^9^-THC also reduced phosphorylation of Akt (aka PKB). Moreover, the CB1R inverse agonist SR 141716A attenuated the negative regulation of neurite outgrowth and ERK1/2 phosphorylation induced by 2AG and Δ^9^-THC. Taken together, our data suggest that hiPSC-derived cortical neurons express CB1Rs and are responsive to both endogenous and exogenous cannabinoids. Thus, hiPSC-neurons may represent a good cellular model for investigating the role of the endocannabinoid system in regulating cellular processes in human neurons.

## Introduction

The endocannabinoid system is a neuromodulatory system with important roles in central nervous system (CNS) development, neuronal function and synaptic plasticity^1,2^. Perturbations in this system have been observed in psychiatric disorders such as schizophrenia^3^, and there is a significant association between cannabis use and schizophrenia^3–6^. The endocannabinoid system is composed of endogenous cannabinoids, enzymes that synthesize and degrade endogenous cannabinoids and cannabinoid receptors. The Cannabinoid 1 receptor (CB1R) is a highly abundant receptor in the CNS, with strong expression in a number of brain regions, including the cortex^1,7^. Cannabinoid 2 receptors (CB2Rs) show much lower expression in the CNS, although recent work has shown high inducible expression of CB2Rs under pathological conditions^8^. Several other receptors, such as peroxisome proliferator activated receptors and transient receptor potential channels, are also engaged by cannabinoids^9^.

CB1Rs are members of the superfamily of G protein-coupled receptors that inhibit adenylyl cyclase and activate mitogen-activated protein kinase by signaling through G_i/o_ proteins. Stimulation of CB1R by cannabinoids has been shown to lead to the phosphorylation of serine/threonine kinase Akt and extracellular signal-regulated kinase ERK1/2^10^. These kinases may then activate or inhibit their substrates to influence cellular functions such as promoting survival, metabolism and differentiation. Both Akt and ERK1/2 activation have been shown to mediate neurite outgrowth^11,12^, a key process in neuronal development and plasticity. Furthermore, the primary psychoactive component of cannabis (*Cannabis sativa*), Δ^9^-tetrahydrocannabinol (Δ^9^-THC) is also thought to act via the CB1R, resulting a wide range of effects including a depression of the glutamatergic system^1^. This highlights the potential importance of the CB1R in relation to health and disease.

Much of our knowledge of the endocannabinoid system comes from work using *in vitro* and *in vivo* animal models. It remains unknown how well these data translate to the human neurons. Recent investigations into the role of endocannabinoids during cortical development have shown that cannabinoids regulate neuronal morphology, through small GTPase signaling pathways and rearrangement of the actin cytoskeleton, resulting in a negative impact on cortical development^13–15^. Recent advances in stem cell technology have enabled the reprogramming of human somatic cells into induced pluripotent stem cells (hiPSCs)^16,17^, which can subsequently be differentiated in to specific neuronal subtypes^18^. Using this system Guennewig and colleagues demonstrated that exposure of hiPSC-neurons to Δ^9^-THC resulted in significant alterations in genes involved with development, synaptic function, as well as those associated with psychiatric disorders^19^. This supports the use of hiPSCs as a cellular model to investigate the cellular actions of cannabinoids to understand their relevance for health and disease.

In this study, we first examined the expression of cannabinoid receptors during the differentiation of hiPSCs into cortical glutamatergic neurons. Subsequently, we investigated the impact of acute exposure to the endogenous cannabinoid 2-arachidonoyl glycerol (2AG), a full agonist for the CB1R, and Δ^9^-THC, a partial agonist for CB1R, on the morphology of young iPSC-derived glutamatergic neurons. Furthermore, we assessed the ability of 2AG and Δ^9^-THC to regulate the activity of ERK1/2, Akt and GSK3β signaling cascades. Finally, we report that the CB1R inverse agonist SR 141716A (Rimonaban), attenuated the negative regulation of neurite outgrowth and ERK1/2 phosphorylation induced by 2AG and Δ^9^-THC. Taken together, our data indicates that CB1R is expressed in immature iPSC-neurons and that 2AG and Δ^9^-THC treatment negatively affect neuronal morphology and impact the Akt and ERK1/2 signaling pathways, potentially via CB1R. These data indicate that cannabinoids may act to influence the development of human cortical neurons.

## Results

### Generation of cortical neurons from hiPSCs

Within the developing CNS, the endocannabinoid system has been implicated in important neuronal processes such as synapse formation and neurogenesis^1,2^. CB1Rs are G-protein coupled receptors that are widely expressed in the brain^7^, including the cortex^1,2^. Consistent with a role for the endocannabinoid system during the development of cortical neurons, acute (24 hour) exposure of hiPSC-neurons to Δ^9^-THC results in the differential expression of multiple genes, including those involved in development^19^. Therefore, in order to further understand the role of cannabinoids during the development of human cortical neurons, we induced cortical differentiation from hiPSC. The cell lines were derived by reprogramming keratinocytes from 3 neurotypic males (age ranging between 35 to 55 years: CTM_01_04, CTM_02_05 and CTM_03_22)^20–23^. HiPSC lines were differentiated into neuroepithelial cell using a dual SMAD inhibition (2i) differentiation protocol for 8 days^20,21,23^ (Figure 1A). The small molecules inhibitors were removed during neuronal progenitor’s differentiation before media was replaced with B27 supplemented with DAPT to induce the generation of terminally differentiated neurons^20,21,23^ (Figure 1A). Immunocytochemical (ICC) analyses indicated that the derived hiPSC expressed high levels of the pluripotency markers OCT4, NANOG and SOX11, whereas early neural progenitor cell (NPC) markers PAX6, ZNF521 and NESTIN were up-regulated following 8 days of 2i induction (Figure 1B and 1C). After 26 days of neuronal differentiation young neurons expressed high levels of FOXG1, TBR1, TBR2 and BRN2 factors, which are required for the differentiation of cortical neurons (Figure 1B and 1C). Similarly, real time PCR (qPCR) analyses demonstrated that the neuroepithelial marker SOX2 was highly expressed 7 days after 2i induction and was gradually down-regulated during the 50 days of neuronal fate acquisition (Figure 1D). PAX6, a key regulator of cortical development, was found to be highly expressed from day 7 to day 50 during neuronal formation; peak expression was observed at day 22 (Figure 1E). The expression of CTIP2, a marker of deep layer cortical cell fate, rapidly increased from day 22 to day 28, peaking at day 26; subsequently, expression of CTIP2 was down-regulated until day 50 (Figure 1F). Conversely, expression of BRN2, a marker associated with upper cortical layer cell fate, showed expression peaks at days 23, 26 and once more at day 48 (Figure 1G). At day 30, no GFAP or S100β positive cells were observed (**Supplementary Figure 1**). ICC analysis at day 50 revealed that most MAP-positive cells were also immunoreactive for markers of glutamatergic fate. This included the -presynaptic proteins synapsin 1 and VGlut1 and the post-synaptic proteins PSD95 and GluN1-subunit of NMDA receptors (Figure 2A-C). To confirm that hiPSC-neurons generated from a hiPSC line derived from keratinocytes, and following our 2i cortical protocol, develop physiological neuronal characteristics, we conducted whole-cell patch clamp recordings. Current clamp recordings demonstrated that these cells were capable of firing an action potential in response to current injection (Figure 2E). Depolarising step potentials (from a holding potential of −60mV) evoked transient inward currents and sustained outward currents, indicative of activation of voltage gated sodium channels and potassium channels respectively (Figure 2E). At this age, we were able to record isolated spontaneous excitatory postsynaptic currents (EPSCs), demonstrating the presence of functional excitatory synaptic connections at this stage (Figure 2F). Collectively, these data demonstrate the generation of cortical glutamatergic neurons from hiPSCs.

**Figure 1:**
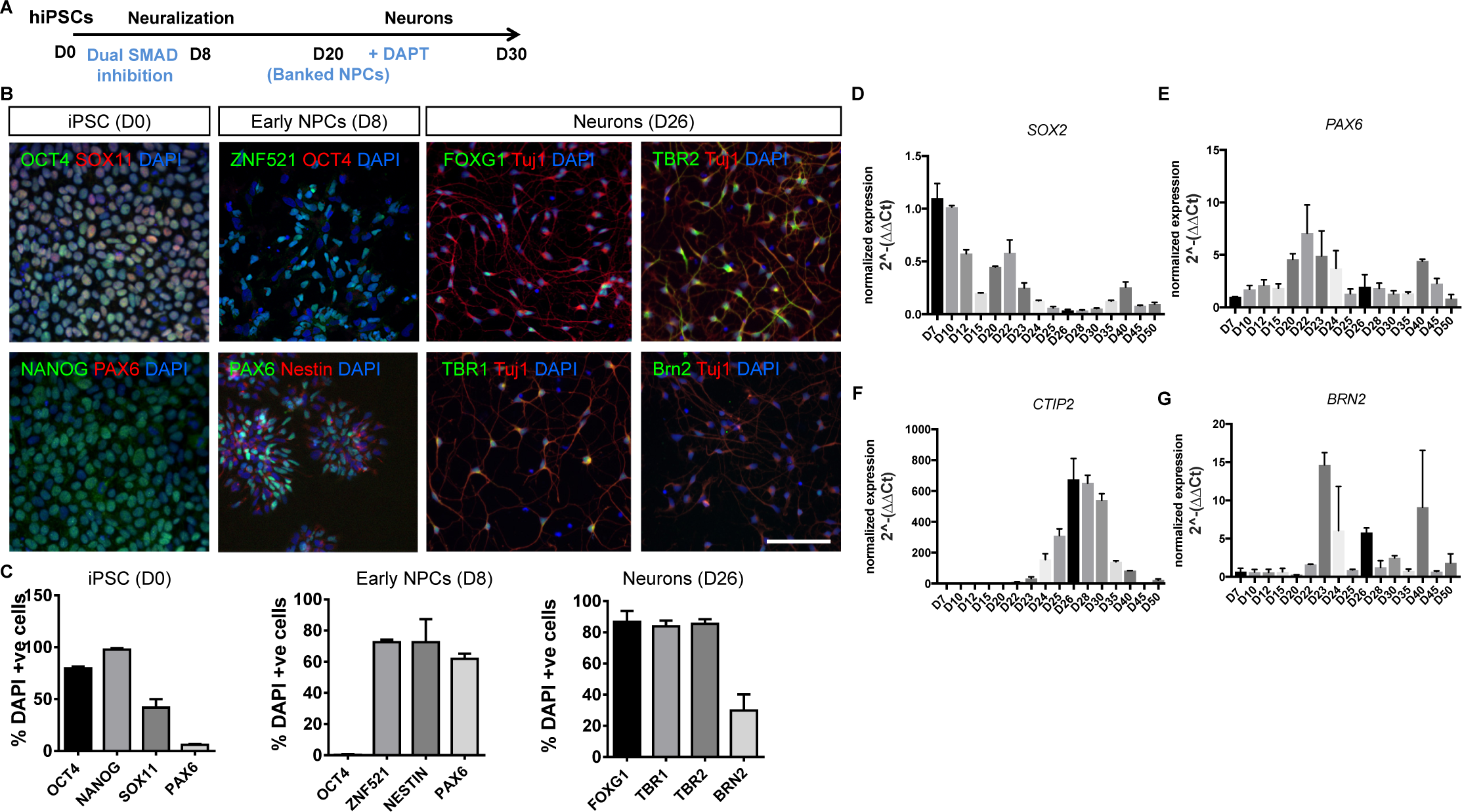
Generation of glutamatergic cortical neurons from hIPSCs. **(A)** Schematic of hiPSC differentiation protocol used. **(B)** Representative images of hiPSCs, NPCs and neurons immunostained for markers of specific cell linages. **(C)** Number of DAPI positive cells expressing markers of hiPSC, NPC and neuronal cell fate; error bars represent standard deviations (SD). **(D-G)** Expression profile of cell fate markers determined between day 7 and 50 following induction of neural differentiation; error bars represent SD.

**Figure 2:**
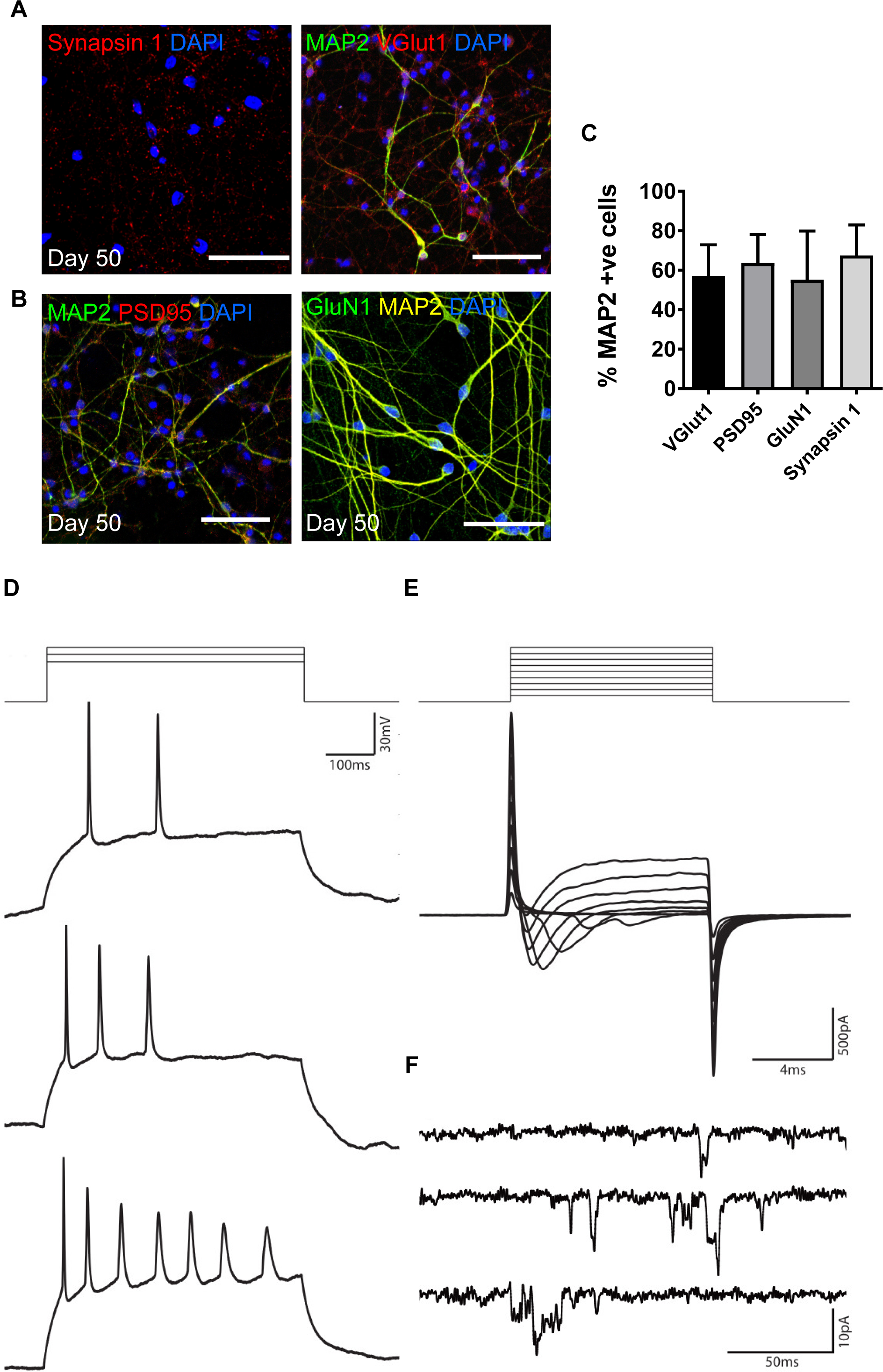
Confirmation of generation of glutamatergic neurons from hiPSCs. **(A & B)** Representative images of day 50 hiPSC-neuron immunostained for glutamatergic markers. **(C)** Number of MAP2 positive cells expressing VGlut1, PSD95, GluN1 and synapsin 1; error bars represent standard deviations (SD). **(D-F)** Whole cell patch clamp recordings obtained from day 64 hiPSC derived neurons. **(D)** Action potentials recorded in response to increasing current injection (24pA, 28pA and 32pA) in current clamp. **(E)** Voltage clamp recordings of the voltage activated currents elicited in response to 10ms long voltage steps from a holding potential of − 70mV. Representative I-V traces at 10mV voltage increments. **(F)** Representative spontaneous current events recorded in voltage clamp at −70mV.

### CB1R expression during the cortical differentiation of iPSCs

Previous studies have suggested that the CB1R is highly expressed during neuronal development^1^. Therefore, we examined the expression of CB1R, at the mRNA level, in hiPSC-derived NPCs and terminally differentiated hiPSC-derived cortical neurons. In all three hiPSC lines, CB1R showed similar gene expression patterns across the three iPSC lines. Specifically, CB1R mRNA levels significantly increased as hiPSCs differentiated from NPCs into neurons (Figure 3A). We also examined the expression of CB2R, GPR55 and TRPV1 receptors which are expressed in the brain and are engaged by cannabinoids in differentiated neurons^1,2^. The mRNA expression levels of these receptors were mirrored across the three hiPSC lines and was significantly reduced compared with the expression of CB1R (Figure 3B). Next, we examined the expression of CB1R in immature hiPSC-neurons using an antibody raised against the C-terminal of the receptor and validated in knockout tissue and human tissue^24,25^ as well by pre-absorption studies (**Supplemental Figure 2A**). Western blotting demonstrated that the receptor was expressed at the protein level in immature hiPSC-neurons. A prominent band ∼53 kDa consistent with the predicated molecular weight for the receptor, was readily observed (Figure 3C). Consistent with these data, immunostaining of immature hiPSC-neurons revealed that CB1R staining could be observed in all three hiPSC clonal lines. CB1R immunoreactivity could be observed within the cell soma, as well as punctate structures along MAP2-positive neurites (Figure 3D). Collectivity, these data demonstrate that CB1R is expressed in immature hiPSC-neurons and is ideally localized to influence neuronal morphology as previously reported.

**Figure 3:**
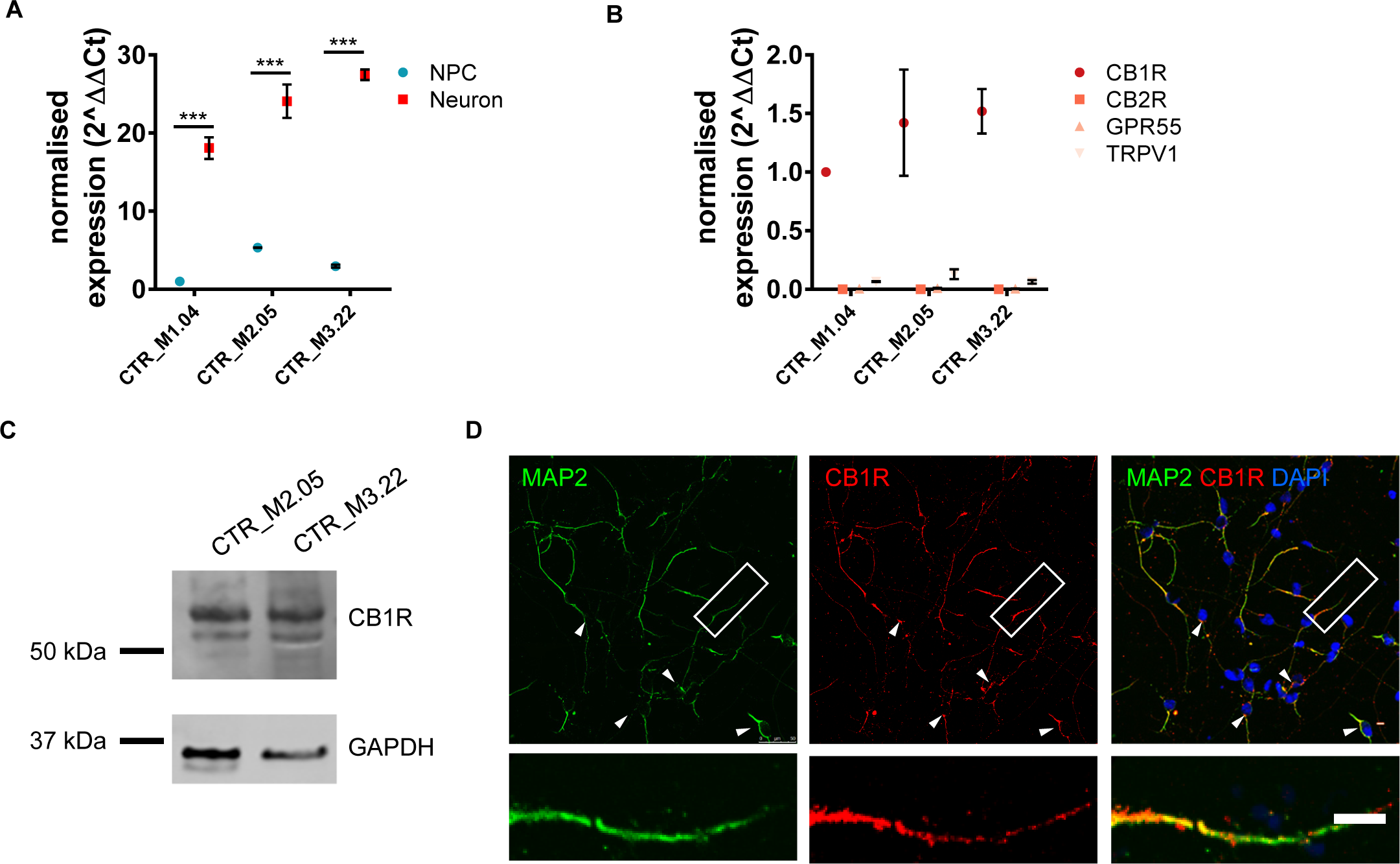
Expression of cannabinoid receptors in hiPSC-neurons. **(A)** Expression of CB1R in NPCs and neurons in three independent hiPSC lines. In all three lines, CB1R expression increases as cells differentiate into neurons: (F(2,12)=6.49, p < 0.05, Bonferroni Post Hoc, ***, p < 0.001, two-way ANOVA; n = 3 independent cultures for each line). **(B)** Comparison of the expression of CB1R, CB2R, GPR55 and TRPV1 receptors in day 30 hiPSC-neurons. Whereas CB1R is highly expressed, no expression of CB2R, GPR55 or TRPV1 receptors was detected in three hiPSC-lines: (F(6,12)=1.896, p < 0.05, Bonferroni Post Hoc, ***, p < 0.001, two-way ANOVA; n = 2 independent cultures for each line). **(C)** Western blot demonstrating expression of CB1R in day 30 hiPSC-neurons derived from 2 independent hiPSC-lines. A prominent band is observed at ∼53 kDa, predicted molecular weight of CB1R. **(D)** Representative confocal images of day 30 hiPSC-neurons immunostained for MAP2 (morphological marker) and CB1R. Immunoreactive puncta for CB1R can be found within the cell soma (white arrow heads) and along MAP2-psoitive neurites. Insets are of high magnification zooms of dendrite highlighted by white box in main image. Scale bar = 5 µm.

### 2AG and Δ^9^-THC negatively regulates neurite outgrowth in hiPSC-neurons

There is growing appreciation that the endocannabinoid system is an important regulator of brain wiring during development through the modulation of several different processes including the specification of neuronal morphology^13,14,26-28^. As CB1R is localized along neurite, we reasoned that endocannabinoids may be involved in regulating the establishment of neuronal morphology in hiPSC-neurons. To test this prediction, we first treated hiPSC-neurons at day 29 with 1 µM 2AG, a full agonist for the CB1R, for 24 hours. Neurons were then fixed and stained for MAP2 to outline neuronal morphology. Neurite outgrowth was assayed using a high-content screening platform (Figure 4A). We first assessed whether treatment with 2AG altered the number of MAP2-positive cells; this revealed no differences between conditions in neurons derived from all 3 hiPSC lines (Figure 4B; p=0.931). We next examined neuronal morphology and demonstrated that the average number of neurites (Figure 4C; p=0.713) and number of branch points (Figure 4C; p=0.780) was unaffected by treatment in neurons derived from all 3 hiPSC lines. However, 2AG treatment for 24 hours significantly reduced neurite length (Figure 4D; F(1,18)=23.4, p < 0.001, Bonferroni Post Hoc, * p < 0.05; two-way ANOVA; n = 3 independent cultures per hiPSC line) in neurons derived from all 3 hiPSC lines.

**Figure 4:**
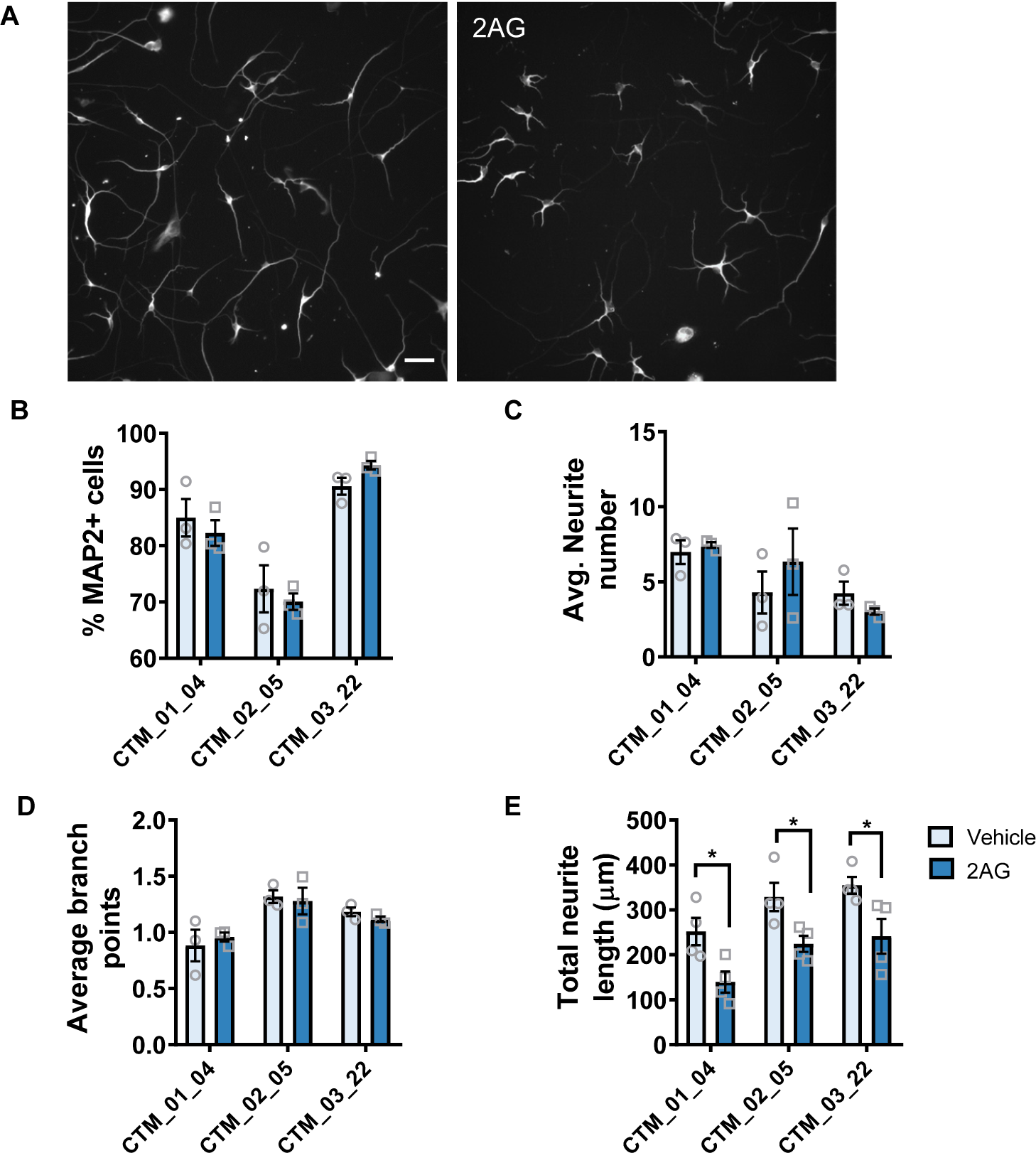
Negative regulation of neurite outgrowth in hiPSC-neurons by 2AG. **(A)** Representative images of MAP2 stained day 30 hiPSC-neurons as imaged using the Opera Phenix High Content Imager. **(B)** Assessing of the number of DAPI-positive MAP2 neurons demonstrates no difference on overall neuronal number across conditions. Data are presented for each individual hiPSC-line: data was generated from 3 independent cultures for each independent hiPSC line. **(C, D)** Assessment of average neurite number **(C)** and average branch number **(D)** revealed no difference between conditions. **(E)** Treatment with 2AG for 24 hours significantly reduced total neurite length compared to vehicle control. Scale bar = 50 µm.

Δ^9^-THC has previously been shown to negatively regulate neurite outgrowth and growth cone dynamics, through the regulation of actin polymerization and microtubule stability in mouse cortical neurons^13,15,26^. In addition, exposure to Δ^9^-THC in hiPSC-neurons results in the alteration of multiple genes involved in the development of neuronal morphology^19^. Therefore, we next examined whether acute exposure to Δ^9^-THC could also alter neurite outgrowth. D29 hiPSC-neurons were treated with 3 µM Δ^9^-THC for 24 hours before neuronal morphology was examined using a high-content screening platform (Figure 5A). Δ^9^-THC treatment did not significantly affect cell number across the three hiPSC-lines (Figure 5B; p=0.647). Similar to 2AG, exposure to Δ^9^-THC did not significantly alter neurite number (Figure 5C; p=0.426) or branch points (Figure 5D; p=0.859). However, Δ^9^-THC treatment for 24 hours significantly reduced the neurite length (Figure 5E; F(1,18)=26.05, p < 0.001, Bonferroni Post Hoc, * p < 0.05; two-way ANOVA; n = 3 independent cultures per hiPSC line) in neurons derived from all 3 hiPSC lines. Taken together, these data indicate that both 2AG and Δ^9^-THC negatively regulate neurite outgrowth in young hiPSC-neurons.

**Figure 5:**
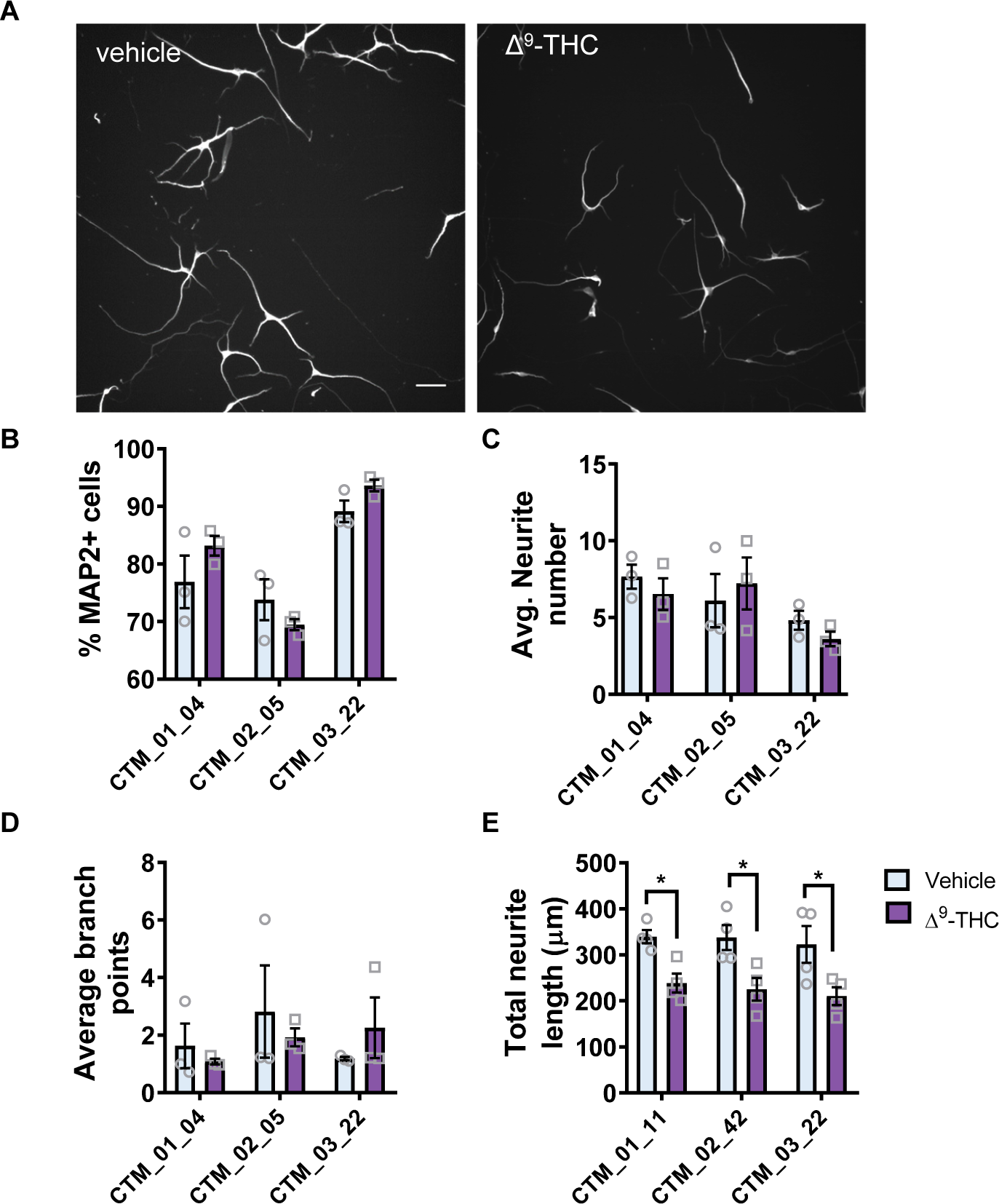
Δ^9^-THC negatively influences neurite outgrowth in hiPSC-neurons. **(A)** Representative images of MAP2 stained day 30 hiPSC-neurons following 24 hour treatment with vehicle or Δ^9^-THC, as imaged using the Opera Phenix High Content Imager. **(B)** Assessing of the number of DAPI-positive MAP2 neurons demonstrates no difference on overall neuronal number across conditions. Data are presented for each individual hiPSC-line: data was generated from 3 independent cultures for each independent hiPSC line. **(C, D)** Assessment of average neurite number **(C)** and average branch number **(D)** revealed no difference between conditions. **(E)** Treatment with Δ^9^-THC for 24 hours significantly reduced total neurite length compared to vehicle control. Scale bar = 50 µm.

### 2AG and Δ^9^-THC modulate phosphorylation of serine/threonine kinases Akt and mitogen-activated protein kinase ERK1/2

Stimulation of CB1R leads to the phosphorylation and activation of several signaling kinases, including Akt, ERk1/2 and GSK3β^2,29,30^. Akt and ERK1/2 pathways have also been implicated in promoting neurite outgrowth^11,12^. GSK-3β, a well-defined substrate of Akt, has also been implicated in the regulation of neurite and axonal outgrowth and branching^31^. Interestingly, the negative effects of CB1R activation on neurite outgrowth have been linked with ERK1/2 signaling^19^, although activation of CB1R has also been linked with Akt/GSK3β signaling *in vivo*^30^. Therefore, we first examined 2AG generated changes in the activation state (phosphorylation) of ERK1/2 and Akt/GSK3β kinases in day 30 hiPSC-neurons. 2AG treatment resulted in a significant decrease in ERK1/2 phosphorylation after 30 minutes (Figure 6A). Conversely, no changes in the phosphorylation state of Akt or GSK3β were detected following 15 and 30 minutes of 2AG exposure (Figure 6B and C). These data demonstrated that 2AG, a full agonist for the CB1R, is capable of negatively regulating ERK1/2 activity.

**Figure 6:**
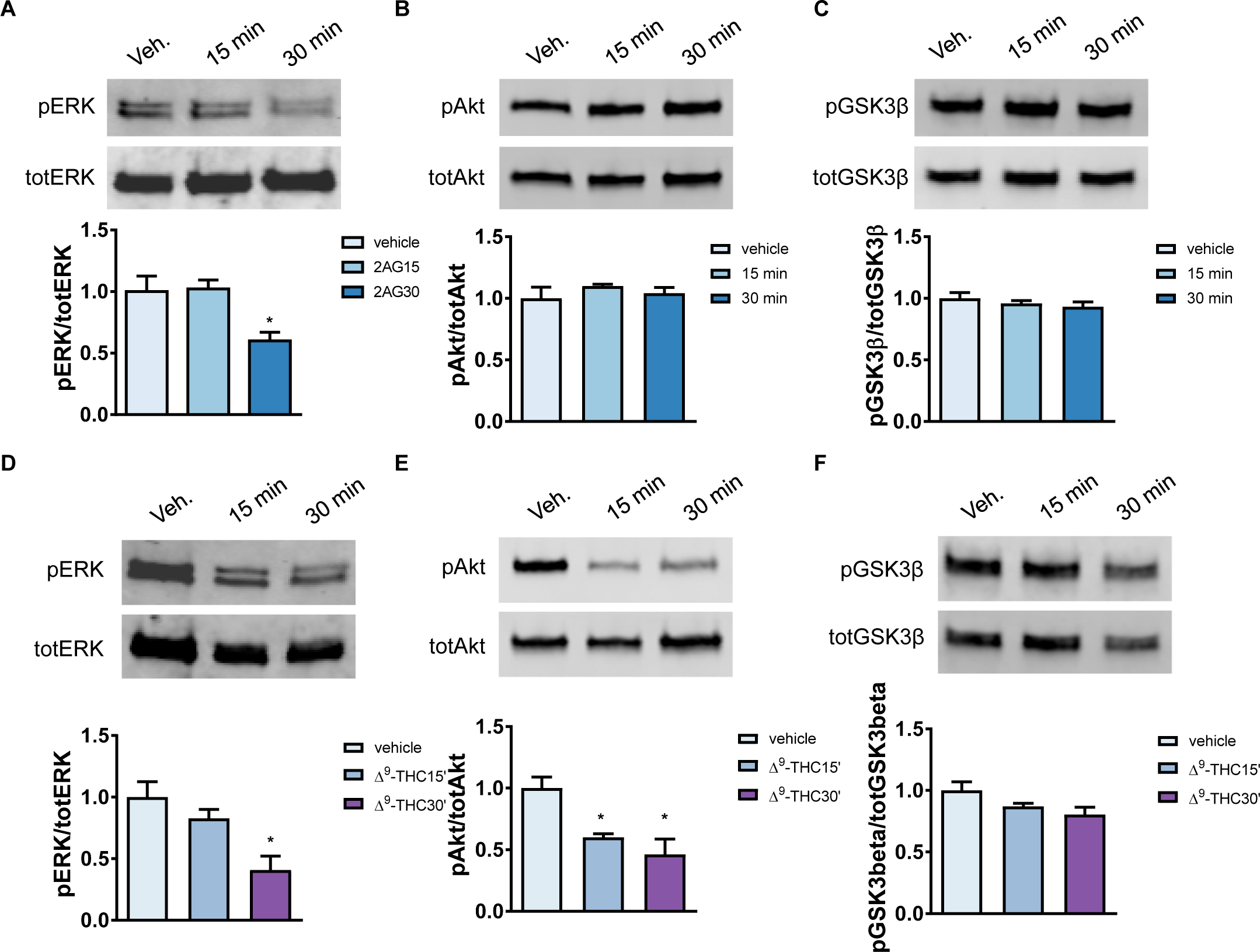
2AG and Δ^9^-THC negative regulate ERK1/2 and Akt phosphorylation. **(A)** Western blot analysis of day 30 hiPSC-neurons treated with 2AG for 15 or 30 minutes compared to vehicle control conditions. Western blots probed for total and pERK1/2, revealed a significant decrease in active (phosphorylated) ERK1/2 levels after 30 minutes of 2AG treatment (F(2,6)=6.593, p < 0.05, Tukey Post Hoc, *, p < 0.05, one-way ANOVA; n = 3 per condition from 3 independent hiPSC-lines). **(B, C)** Cell lysates were also assessed for levels of active (phosphorylated) Akt and GSK3β; no difference was observed between conditions. **(D)** Western blot analysis of day 30 hiPSC-neurons treated with Δ^9^-THC for 15 or 30 minutes compared to vehicle control conditions. Western blots probed for total and pERK1/2, revealed a significant decrease in active (phosphorylated) ERK1/2 levels after 30 minutes of Δ^9^-THC treatment (F(2,6)=8.299, p < 0.05, Tukey Post Hoc, *, p < 0.05, one-way ANOVA; n = 3 per condition from 3 independent hiPSC-lines). **(E)** Assessment of pAkt levels also revealed a reduction in active levels of Akt following treatment with Δ^9^-THC (F(2,6)=9.557, p < 0.05, Tukey Post Hoc, *, p < 0.05, one-way ANOVA; n = 3 per condition from 3 independent hiPSC-lines). **(F)** Cell lysates were also assessed for levels of active (phosphorylated) GSK3β; no difference was observed between conditions.

Owing to our data demonstrating that 2AG negatively regulated ERK1/2 phosphorylation, we next investigated the effect of Δ^9^-THC on ERK1/2, Akt and GSK3β kinases. D30 hiPSC-neurons were treated with vehicle or 3 µM Δ^9^-THC for either 15 or 30 minutes. Similar to 2AG treatment, Δ^9^-THC induced a rapid decrease in ERK1/2 phosphorylation (Figure 6D). Interestingly, Δ^9^-THC also caused a rapid reduction in levels of phosphorylated Akt, which was evident after 15 minutes (Figure 6E). A non-significant trend towards a decrease in phosphorylated GSK3β was also evident following Δ^9^-THC treatment, but this did not reach significance (Figure 6F; p=0.105). Collectively, these data indicate that Δ^9^-THC negatively regulates both ERK1/2 and Akt phosphorylation, in contrast to 2AG, which regulates ERK1/2 exclusively.

### 2AG and Δ^9^-THC modulate Akt and ERK1/2 phosphorylation via the CB1R receptor

As our data indicated that both 2AG and Δ^9^-THC negatively regulated ERK1/2 and Akt signaling kinases, we were interested in understanding whether these effects were being mediated by the CB1R. Recently, Obiorah et al. showed that Δ^9^-THC mediated changes in glutamate receptor gene expression in hiPSC-neurons could be blocked using the CB1R selective inverse antagonist SR 141716A (Rimonabant)^32^. Pre-treatment with 20 nM SR 141716A blocked 2AG and Δ^9^-THC (30 minutes) induced de-phosphorylation of ERK1/2; SR 141716A alone had not effect on pERK1/2 levels (Figure 7A; F(1.309,2.618)=12.07, p = 0.0489, Tukey Post Hoc, * p < 0.05; one-way ANOVA). As previously seen, Δ^9^-THC, but not 2AG (30 minutes) caused a reduction in pAkt levels. This effect was blocked by pre-treatment with 20 nM SR 141716A (Figure 7B; F(1.542,3.083)=14.15, p = 0.0291, Tukey Post Hoc, * p < 0.05; one-way ANOVA). Neither 2AG or Δ^9^-THC (30 minutes) had an effect on GSK3β; SR 141716A alone had no effect on the activity of this kinase (Figure 7C; F(1.747,3.494)=0.9429, p = 0.4567; one-way ANOVA). Taken together, these data indicate that 2AG and Δ^9^-THC negatively regulated ERK1/2 and Akt signaling kinases by acting through the CB1R receptor.

**Figure 7:**
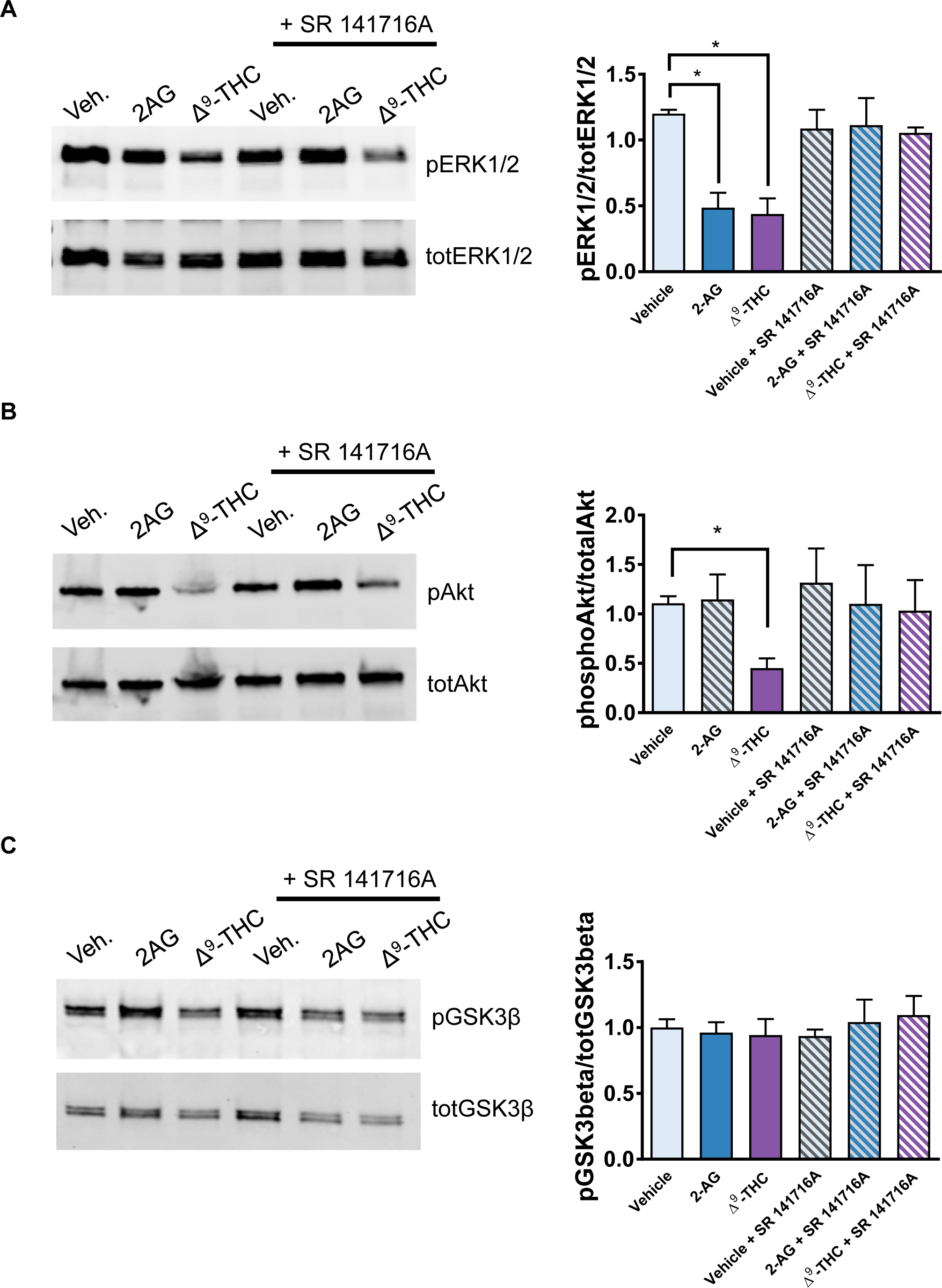
CB1R antagonist SR 141716A blocks negative regulation of ERK1/2 and Akt phosphorylation by 2AG and Δ^9^-THC. **(A)** Western blot analysis of day 30 hiPSC-neurons treated with vehicle, 2AG or Δ^9^-THC (30 minutes) with or without pre-treatment with inverse antagonist SR 141716A (20 nM). Western blots probed for total and pERK1/2, revealed that SR 141716A blocked 2AG or Δ^9^-THC induced decrease in active (phosphorylated) ERK1/2 levels after 30 minutes. **(B)** Cell lysates were also assessed for levels of pAkt. This revealed that SR 141716A blocked Δ^9^-THC decrease in pAkt levels. **(C)** Cell lysates were also assessed for levels of active (phosphorylated) GSK3β; no difference was observed between conditions.

**Figure 8:**
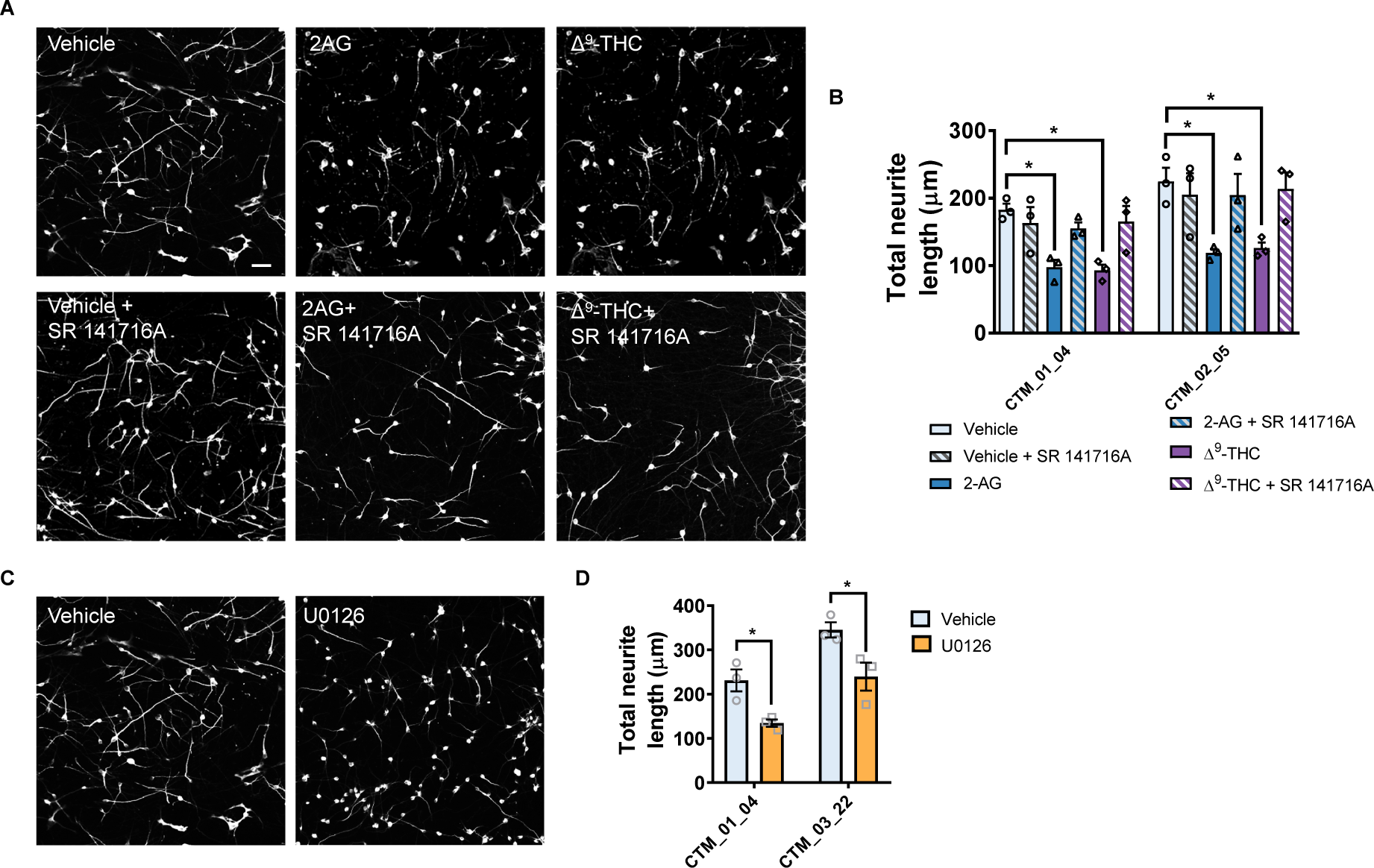
Negative regulation of neurite outgrowth by 2AG and Δ^9^-THC is attenuated by SR 141716A. **(A)** Representative images of MAP2 stained day 30 hiPSC-neurons following 24 hour treatment with vehicle, 2AG or Δ^9^-THC, with or without co-treatment with inverse antagonist SR 141716A (20 nM). **(B)** Assessing of total neurite length in all conditions. Data are presented for each individual hiPSC-line: data was generated from 3 independent cultures for each independent hiPSC line. Treatment with 2AG or Δ^9^-THC for 24 hours significantly reduced total neurite length compared to vehicle control. This effect was attenuated by SR 141716A. **(C)** Representative images of MAP2 stained day 30 hiPSC-neurons following 24 hour treatment with vehicle or U0126 (10 μM). **(D)** Assessing of total neurite length in all conditions. Data are presented for each individual hiPSC-line: data was generated from 3 independent cultures for each independent hiPSC line. Treatment with U026 significantly reduced total neurite length compared to vehicle control. Scale bar = 50 µm. Western blot analysis of day 30 hiPSC-neurons treated with vehicle, 2AG or Δ^9^-THC (30 minutes) with or without pre-treatment with inverse antagonist SR 141716A. Western blots probed for total and pERK1/2, revealed that SR 141716A blocked 2AG or Δ^9^-THC induced decrease in active (phosphorylated) ERK1/2 levels after 30 minutes. **(B)** Cell lysates were also assessed for levels of pAkt. This revealed that SR 141716A blocked Δ^9^-THC decrease in pAkt levels. **(C)** Cell lysates were also assessed for levels of active (phosphorylated) GSK3β; no difference was observed between conditions.

### Negative regulation of neurite outgrowth by 2AG and Δ^9^-THC is blocked by CB1R antagonism

As our data indicated that the CB1R mediates the effects of 2AG and Δ^9^-THC on ERK1/2 activity, we next questioned whether this receptor was required for the negative regulation of neurite outgrowth y these cannabinoids. To test this, we treated day 29 hiPSC-neurons with 2AG or Δ^9^-THC with or without SR 141716A. Treatment with 2AG or Δ^9^-THC caused a reduction in total neurite length; this was attenuated by co-treatment with SR 141716A (Figure 7A and B; F(1,24)=12.25, p=0.0018, Bonferroni Post Hoc, * p < 0.05; two-way ANOVA; n = 3 independent cultures per hiPSC line). Next, we reasoned that if 2AG or Δ^9^-THC were causing a reduction in neurite outgrowth by negatively regulating the ERK1/2 pathway, that inhibiting this kinase pathway would mimic the effects of these cannabinoids on neurite outgrowth. Therefore, we treated day 29 hiPSC-neurons with 10 μM U0126, a MEK inhibitor. Assessment of neurite outgrowth on day 30 revealed that neurons treated with U0126 had reduced total neurite length (Figure 7A and B; F(1,8)=24.1, p=0.0012, Bonferroni Post Hoc, * p < 0.05; two-way ANOVA; n = 3 independent cultures per hiPSC line). Taken together, these data indicate that antagonism of the CB1R attenuates the negative effect on neurite outgrowth induced by 2AG and Δ^9^-THC. Furthermore, inhibiting the MEK/ERK1/2 pathway mimics the effect of these cannabinoids on neurite outgrowth.

## Discussion

Recently it has been demonstrated that cannabinoid treatment affects neuronal function and glutamate receptor expression in hiPSC-derived dopaminergic neurons and excitatory neurons, respectively^32,33^. Moreover, treatment with Δ^9^-THC induces a wide range of transcriptional changes in cortical hiPSC-neurons. This includes changes in the expression of genes involved in neurodevelopmental processes, as well as those associated with neurodevelopmental and psychiatric disorders^19^. Despite this, it is not clear what role the endocannabinoid system plays during neurodevelopment. In this study, we demonstrate that the CB1R is the major cannabinoid receptor expressed during early differentiation of cortical hiPSC-neurons. Furthermore, we demonstrate that treatment with both 2AG and Δ^9^-THC result in a negative regulation of neurite outgrowth, as well as differential effects on the activation status of ERK1/2 and Akt signaling kinases. Furthermore, out data indicate that both 2AG and Δ^9^-THC are likely signaling via the CB1R receptor to induce these effects, but are able to induce different signaling patterns, potentially owing to their pharmacological profile at these receptors.

Differentiated hiPSC-derived glutamatergic neurons showed significant upregulation of CB1R mRNA compared to undifferentiated NPCs. Moreover, hiPSC-derived glutamatergic-neurons also showed significantly higher levels of CB1R mRNA compared to CB2R, GPR55 and TRPV1. Stanslowsky et al. (2017) also observed upregulation of CB1R mRNA following terminal differentiation of hiPSC-derived dopaminergic neurons and the predominant expression of CB1R in hiPSC-derived neural precursors^33^. It has been reported that CB1R is highly expressed in newly differentiated glutamatergic neurons and is labelled in a punctate manner in the soma and neurites of these cells^26^. In agreement with previous reports, we observed that CB1R mRNA is differentially up-regulated during neuronal differentiation in the three iPSC lines with different genetic background. Moreover, at the protein level, CB1R was expressed in immature hiPSC-derived glutamatergic neurons, were it localized in the cell soma and along neurites. These findings provide support for the idea that the CB1R may play an important role in early neurodevelopmental processes such as progenitor cell differentiation and neurite outgrowth, prior to the formation of synapses.

Studies on the effect of cannabinoids on neurite outgrowth have been attempted with various CB1R agonists and have produced conflicting results^2^. Whereas Δ^9^-THC treatment in murine dopaminergic neurons had no effect on morphology^34^, Δ^9^-THC treatment reduced neurite length in primary neurons from E16.5 mouse cortices^15^. Similarly, activation of CB1R by HU210 has been shown to trigger neurite outgrowth in Neuro2A cells^35^, while CB1R agonist WIN55,212-2 negatively regulates dendritic and axonal outgrowth in cultured rat hippocampal neurons^26^. We investigated the effects of the endogenous cannabinoid 2AG and the exogenous cannabinoid Δ^9^-THC on neuronal morphology. In our study, we observed that both 2AG and Δ^9^-THC decreased total neurite outgrowth in hiPSC-derived glutamatergic neurons. Vitalis et al. (2008) previously showed that basal activation of CB1R acts as a negative regulatory signal for neuritogenesis, and treatment with inverse agonist AM281 blocked these effects^26^. These data and in addition to those presented in this study suggest that both endogenous and exogenous cannabinoids affect neuronal morphology in growing neurons. However, the specific downstream mechanisms underlying this process are not well-described, and further research is required to delineate how different cannabinoids affect neuritogenesis in growing and mature neurons, from humans and other animal species.

CB1R signaling is a complex process and cannabinoids can phosphorylate and activate members of all three families of multifunctional mitogen-activated protein kinases, as well as the PI3K/Akt pathway^36,37^. We examined the signal transduction pathways regulated by the activation of CB1Rs, focusing on the effect of 2AG and Δ^9^-THC on the Akt and ERK1/2 pathways. Studies on the effect of cannabinoids on these pathways have been attempted with various CB1R agonists and have produced conflicting results. Several studies of non-neuronal cells and hippocampal slices and hippocampal neurons in living mice previously demonstrated that stimulation of CB1Rs by endocannabinoids anandamide, 2AG and exogenous Δ^9^-THC activates ERK1/2^29,38^, whereas stimulation of CB1R by selective CB1R agonist ACEA in primary cortical neurons did not alter the phosphorylation of ERK1/2 or Akt ^39^. In contrast, an increase in the phosphorylation of Akt and of GSK3β by acute Δ^9^-THC, anandamide and HU-210 administration has been demonstrated *in vivo*^30,40^. In this study, 2AG did not affect the phosphorylation and activation of Akt, whereas Δ^9^-THC reduced the levels of phosphorylated Akt. Neither cannabinoid significantly altered the downstream effector GSK-3β, though both reduced the levels of phosphorylated ERK1/2. Overall, previous data and ours suggest that both endogenous and exogenous cannabinoids affect the ERK1/2 and Akt signaling pathways. It is possible that in different model systems, CB1R activation triggers different signaling pathways to regulate downstream gene expression and neuronal processes. The specific pharmacology of CB1R signaling is not well understood, and further research is necessary to describe how endogenous, exogenous and synthetic cannabinoids act on CB1R and how they may or may not interact to trigger or inhibit downstream signaling pathways.

Overall, the results of the present study suggest that young hiPSC-derived glutamatergic neurons are responsive to cannabinoids, and further support the notion that this cellular system is a good model for investigating CB1R signaling^19^. CB1Rs were first identified as the main neuronal receptor for Δ^9^-THC and are one of the most abundant G protein-coupled receptors in the brain^7^. These receptors have previously been characterized in being involved in retrograde synaptic signaling, acting to control neuronal activity in mature neurons^41–43^ and to affect both long-term potentiation (LTP) and depression (LTD) and to impair learning and memory^44^. CB1Rs and endocannabinoids are also highly expressed in the fetal brain and implicated in neuronal development processes such as neurite growth and axonal pathfinding^13,15,26–28,35^. As shown in this and other recent studies, both endogenous and exogenous cannabinoids alter neurite outgrowth, synaptic activity and synaptic protein expression in developing human hiPSC-derived neurons^32,33^. Moreover, as Δ^9^-THC has recently been shown to alter expression of genes associated with neurodevelopmental and psychiatric disorders, there is evidence for perturbations of shared molecular pathways potentially exacerbated by Δ^9^-THC^19^. Thus, these data collectively suggest that cannabinoids may play a role in brain development. Furthermore, they raise the possibility that overstimulation by exogenous cannabinoids or abnormal levels of endocannabinoids may perturb normal physiological processes, during neurodevelopment which may contribute to the pathophysiology of neurodevelopmental and psychiatric disorders, however further in-depth studies are required to investigate these possibilities.

## Materials and Methods

### Reagents

Δ^9^-THC (ethanol solution, ab120447) were from Abcam. 2-Arachidonylglycerol (1298), U0126 (1144) and SR 141716A (0923) were from Tocris. Δ^9^-THC was prepared by serially diluting the solution in culture media to a 10x stock solution. The CB1R inverse antagonist SR 141716A was dissolved in DMSO and used at a final concentration of 20 nm; an equivalent volume of DMSO or ethanol was used as a vehicle control. A list of antibodies used in this study can be found in **Table 1**.

### Human induced pluripotent stem cells (hiPSCs)

hiPSC lines were generated from primary keratinocytes as described previously^20^. Participants were recruited and methods carried out in accordance to the ‘Patient iPSCs for Neurodevelopmental Disorders (PiNDs) study’ (REC No 13/LO/1218). Informed consent was obtained from all subjects for participation in the PiNDs study. Ethical approval for the PiNDs study was provided by the NHS Research Ethics Committee at the South London and Maudsley (SLaM) NHS R&D Office. Briefly, 1 x 10^5^ primary hair root keratinocytes were reprogramed by introducing OCT4, SOX2, KLF4 and C-MYC factors with a CytoTune-iPS 2.0 Sendai expressing Reprogramming Kit (ThermoFisher, A16517). Transformed keratinocytes were plated onto an irradiated MEF feeder layer (Millipore) supplemented Epilife medium for ten days before switching to ‘hES media’, which consisted of KO-DMEM/F12 supplemented with 20% Knock-out serum replacement, Non-essential amino acids, Glutamax, β-mercaptoethanol (all from Life Technologies) and bFGF (10 ng/ml; Peprotech). After a further two weeks, reprogrammed colonies were selected and plated on Geltrex (Life technologies) coated Nunc treated multidishes (Thermo Scientific) into E8 media (Life Technologies). hiPSCs reprograming was validated by genome-wide expression profiling using Illumina Beadchip v4 and the bioinformatics tool ‘Pluritest’. Additionally, the tri-lineage differentiation potential was stablished by embryoid body formation; ICCs to validate the expression of different pluripotency markers including Nanog, OCT4, SSEA4 and TRA1-81 and the alkaline phosphatase activity by Alkaline phosphatase expression kit (Milipore). The genomic stability was determined by G-banded karyotyping. hiPSCs were incubated in hypoxic conditions at 37°C and maintained in E8 media replaced every 24 hours until the cells monolayer reach ∼95% confluence.

### Neuronal differentiation

Neuronal differentiation of hiPSCs was by achieved replacing E8 medium on confluent hiPSCs with neuralization medium: 1:1 mixture of N2- and B27 (Life Technologies) supplemented with 10 μM SB431542 (Sigma-Aldrich) and 1 μM Dorsomorpin (Calbiochem) for dual SMAD inhibition (2i). Cells were maintained at 37°C in normoxic conditions in 2i medium for 7 days with media was replacement every 24 hours. At day 7 the 2i induced neuroepithelial cells were then passaged using Accutase (Life Technologies) and re-plated at a 1:1 ratio in N2:B27 media without the small molecule inhibitors and supplemented with 10 μM Rock Inhibitor Y-27632 (Sigma Aldrich) until day 12. Cells were then passaged 2 more times following the same procedure described above at days 16 and 20. During neuronal induction, the formation of neural rosettes was evident from approximately day 10 until the differentiation of neural progenitors (NPC) around day 20. NPCs were maintained as mitotic progenitors in neuralization medium supplemented with 10 ng/ml bFGF, 5 µg/ml insulin, 1 mM l-glutamine, 100 µm non-essential amino acids, and 100 µM 2-mercaptoethanol. Dividing cells were routinely passaged to a 1:1 ratio to expand the NPC population as necessary. Moreover, NPC vials were frozen in a controlled freezing rate using Mr. Frosty containers in 10% DMSO in NPC media (Thermo Fisher). NPC cryovials were thaw by incubation for 3 minutes at 37°C and then transferring the cell suspension to a 15 ml tube containing 3 ml of NPC media. Subsequently, the tubes were centrifuged (1250RPM for 2 minutes) and the cells pellet was re-suspended with 3 ml of NPC + 10 µM Rock inhibitor media and plated as above. NPC were terminally plated on 5 µg/ml poly-D-lysine and 2 µg/cm^2^ laminin (Thermo Fisher) in coated Nunc treated multidishes, in B27 medium supplemented with 200 µM L-ascorbic acid and 10 µM DAPT (Calbiochem) for 7 days (day 27) to block NOTCH signaling. Subsequently, neurons were grown in B27 medium with 200 µM L-ascorbic acid until day 30 when they were used for experimentation.

### Treatments

Acute pharmacological treatment (15 or 30 minutes) of cells were performed in ACSF solution (125 mM NaCl, 2.5 mM KCl, 26.2 mM NaHCO_3_, 20 mM glucose, 5 mM Hepes, 2.5 mM CaCl_2_ and 1.25 mM MgCl_2_). Prolonged pharmacological treatment (24 hour) were performed in Neurobasal medium supplemented with B27 and 200 µM L-ascorbic acid.

### qRT-PCR

Total RNA harvested and lysed with Trizol reagent (Life technologies) and isolated by centrifugation with 100% Chloroform, following by 100% isopropanol and lastly by 75% ethanol. The RNA was purified by precipitation with 100% ethanol and Sodium acetate (Life technologies) and quantify with the NanoDrop 1000 Spectrophotometer (Thermo scientific). Residual genomic DNA was removed by addition of TURBO DNA-free (Life technologies) and incubation at 37°C for 30 minutes. Complementary DNA (cDNA) was synthesized from 1 µg of total RNA from each extraction using random hexamer primers and SuperScript III (Life Technologies) following the manufacturer’s recommendations. qPCR was performed with HOT FIREPol EvaGreen qPCR Mix Plus ROX (Solis Biodyne) carried out according to the manufacturer’s instructions in a total volume of 20 µl, containing 1:5 diluted cDNA, qPCR mix and primers at to a final concentration of 0.3 μM. PCR reaction conditions: 95°C for 15 minutes for the initial denaturation followed by 95°C for 30 seconds, 60°C for 30 seconds and 72°C for 30 seconds during 33 cycles. The melting curve analyses was preformed from 60°C to 95°C with readings every 1°C. The 2^ Δ Δ CT comparative method for relative quantification was used to quantify the genes expression. The data CT values were normalized to GAPDH, RPL27 and SDHA housekeeping genes.

### Immunocytochemistry

Treated neurons were fixed with 4% formaldehyde plus 4% sucrose in PBS. Fixed neurons were permeabilized in 0.1% Triton-X-100 in PBS for 15 minutes and blocked in 4% normal goat serum in PBS for 1 hour at room temperature. Primary antibodies were added to the block solution in an antibody dependent concentration and incubated overnight at 4°C. Immunoreactivity was achieved by incubating the cells with 1:500 concentration of Alexa Fluor 594 conjugated anti-mouse IgG, Alexa Fluor 594 conjugated anti-goat IgG and Alexa Fluor 488 conjugated anti-rabbit IgG in block buffer. For nuclei staining a 1:2000 concentration of DAPI (Thermo Fisher) was used.

### Imaging of immunofluorescence by high content image screening

NPCs were plated at a density of 1×10^4^ cells/well on poly-D-lysine and laminin-coated optical-bottom 96 well plates with polymer base (ThermoScientific). Image acquisition was performed with a 20X objective for the genes OCT4, NANOG, SOX11, ZNF521, PAX6, NESTIN, FOXG1, TBR1, TBR2 and BRN1 by using CellInsight CX5 High Content Screen Platform (Thermo Fisher). The bioapplication Cell Health Profiling from the iDev software package (Thermo Fisher) using the nuclear staining to assessed the viable cells. Intensity, shape and size features were used to determine positively labelled cells.

For neurite outgrowth assays, treated hiPSC-neurons were imaged using an Opera Phenix High Content screening platform (Perkin Elmer): images were acquired using a 20 x (NA 0.4) objective. The Harmony High Content Imaging and Analysis Software was used to determine the average neurite length and average branch number based on the MAP2 staining. For each hiPSC line, 3 biological replicates with 3 technical replicates per condition were imaged and analyzed: 15 randomly selected fields from each technical replicate was examined. Data from each technical replicate was averaged and used as a single data point and compared with each biological replicate and each hiPSC line. The means of percentages of positive cells and the average neurite length and branch number were compared by an ANOVA. Confocal images of hiPSC-neurons stained for MAP2 and CB1R were acquired using a Lecia SP5 confocal microscope using a 63x oil objective (NA 1.4) as a z-series. Post image acquisition, images were z-projected using ImageJ (rsb.info.nih.gov/ij/).

### Western blotting

hiPSC-neurons cells were lysed in 20 mM Tris, pH 7.2, 150 mM NaCl, 1% Triton-X-100, 5 mM EDTA pH 8 containing a cocktail of protease and phosphatase inhibitors. Detergent soluble lysate were resolved by SDS-PAGE, then immunoblotted with primary antibodies overnight at 4°C, followed by incubation with anti-mouse Alexa Fluor 680 and anti-rabbit Alexa Fluor 790 IgG (H+L) secondary antibodies for 1 hour at room temperature. Membranes were scanned with Odyssey imaging system (LI-COR). Intensities of bands were quantified by densitometry using Image J (rsb.info.nih.gov/ij/). Full length blots are shown in **Supplemental Figures 1-3.**

### Electrophysiology

Electrophysiological recordings were obtained from hiPSC derived neurons differentiated for 64 days *in vitro*. Patch pipettes (4.0–7.5 MΩ) were pulled from borosilicate glass capillary tubes using a P97 Flaming/Brown Micropipette Puller (Sutter Instruments). The internal patch solution contained (in mM) 135 K-Gluconate, 10 KCl, 1 MgCl2, 10 HEPES, 2 Na2-ATP and 0.4 Na3-GTP. All recordings were conducted at room temperature in neurobasal medium (0.1% BDNF, 0.01% Ascorbic acid, 2% B27, 1% Glutamax). Spontaneous currents were measured by whole cell voltage clamp recordings at a holding potential of −70mV. Current-voltage recordings were conducted from a holding potential of −70mV with increasing voltage steps of 2mV increments up to +20mV. Action potentials were recorded in current clamp from a holding potential of −6 mV, with 2pA current injection increments each of 500ms. Data were generated and acquired using an EPC10 amplifier (Heka Instruments, Bellmore, NY, USA) and the software PatchMaster.

### Statistical analysis

All statistical analysis was performed in GraphPad. Differences in 2^ Δ Δ CT, relative expression, cell number and neurite parameters were identified by Student’s unpaired t-tests, or for comparisons between multiple conditions the main effects and simple effects were probed by one-way-ANOVAs or two-way-ANOVAs with Tukey or Bonferroni correction for multiple comparisons. Differences were considered significant if *P* was lower than 0.05 (p < 0.05). Error bars represent standard errors of the mean unless stated otherwise.

## Acknowledgements

The study was supported by grants from the Wellcome Trust ISSF Grant (No. 097819) and the King’s Health Partners Research and Development Challenge Fund, a fund administered on behalf of King’s Health Partners by Guy’s and St Thomas’ Charity awarded to DPS; the Brain and Behavior Foundation (formally National Alliance for Research on Schizophrenia and Depression (NARSAD); Grant No. 25957), awarded to DPS; the Innovative Medicines Initiative Joint Undertaking under grant agreement no. 115300, resources of which are composed of financial contribution from the European Union’s Seventh Framework Programme (FP7/2007-2013) and EFPIA companies’ in kind contribution (JP and DPS). This work was also supported by funds the European Autism Interventions (EU-AIMS), and the Innovative Medicines Initiative Joint Undertaking under grant agreement no. 115300, resources of which are composed of financial contribution from the European Union’s Seventh Framework Programme (FP7/2007-2013) and EFPIA companies’ in kind contribution (JP, DPS and LCA); a Medical Research Council 4-year PhD studentship to SET, and funds from the Mortimer D Sackler Foundation and the Sackler Institute for Translational Neurodevelopment (RDT). SB has received support from the NIHR (NIHR Clinician Scientist Award; NIHR CS-11-001) and the UK MRC (MR/J012149/1). We thank the Wohl Cellular Imaging Centre (WCIC) at the IoPPN, Kings College, London, for help with microscopy.

## Author contributions

C.S., L.D., E.A., K.W.C., S.E.T., R.D.T., L.C.A and D.P.S. carried out experiments and data analysis; N.J.B., J.P. and S.B. provided reagents, assistance in experimental design and editing of manuscript; C.S., L.D. and D.P.S. wrote the manuscript; D.P.S. designed the project.

