## Supplemental Figures for "∆^9^-tetrahydrocannabinol negatively regulates neurite outgrowth and Akt signaling in hiPSC-derived cortical neurons"

^1^Department of Basic and Clinical Neuroscience, The Maurice Wohl Clinical Neuroscience Institute, Institute of Psychiatry Psychology and Neuroscience, King's College London, London, SE5 8AF, UK; ^2^MRC Centre for Neurodevelopmental Disorders, King’s College London, London, UK; ^3^Centre for Developmental Neurobiology, King's College London, London, UK, ^4^Department of Psychiatry, University of Oxford, UK; ^5^National Institute for Biological Standards and Control, South Mimms, UK ^6^Department of Psychosis Studies, King’s College London, London, SE5 8AF, UK.

**Supplementary Information:**

**Supplementary Figures 1-5**

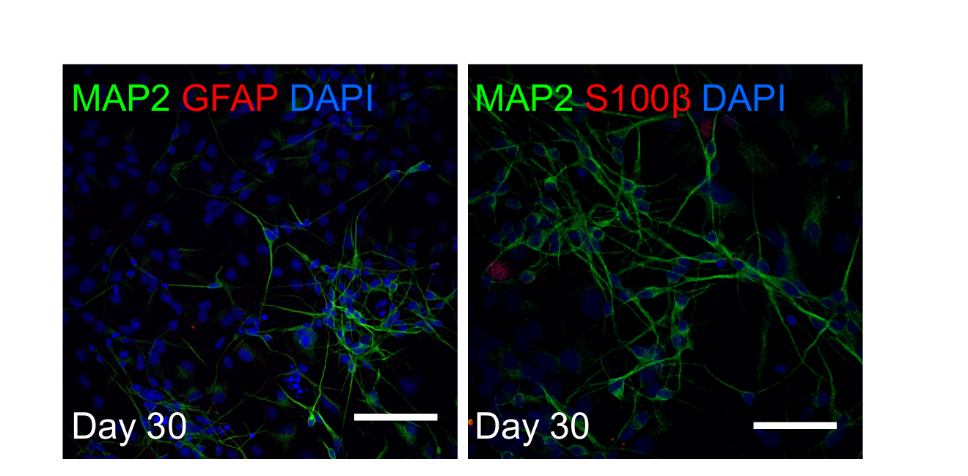

**Supplementary Figure 1.** Representative images of day 30 hiPSC-neurons immunostained for glia markers GFAP and S100β. Scale bar = 50 μM.

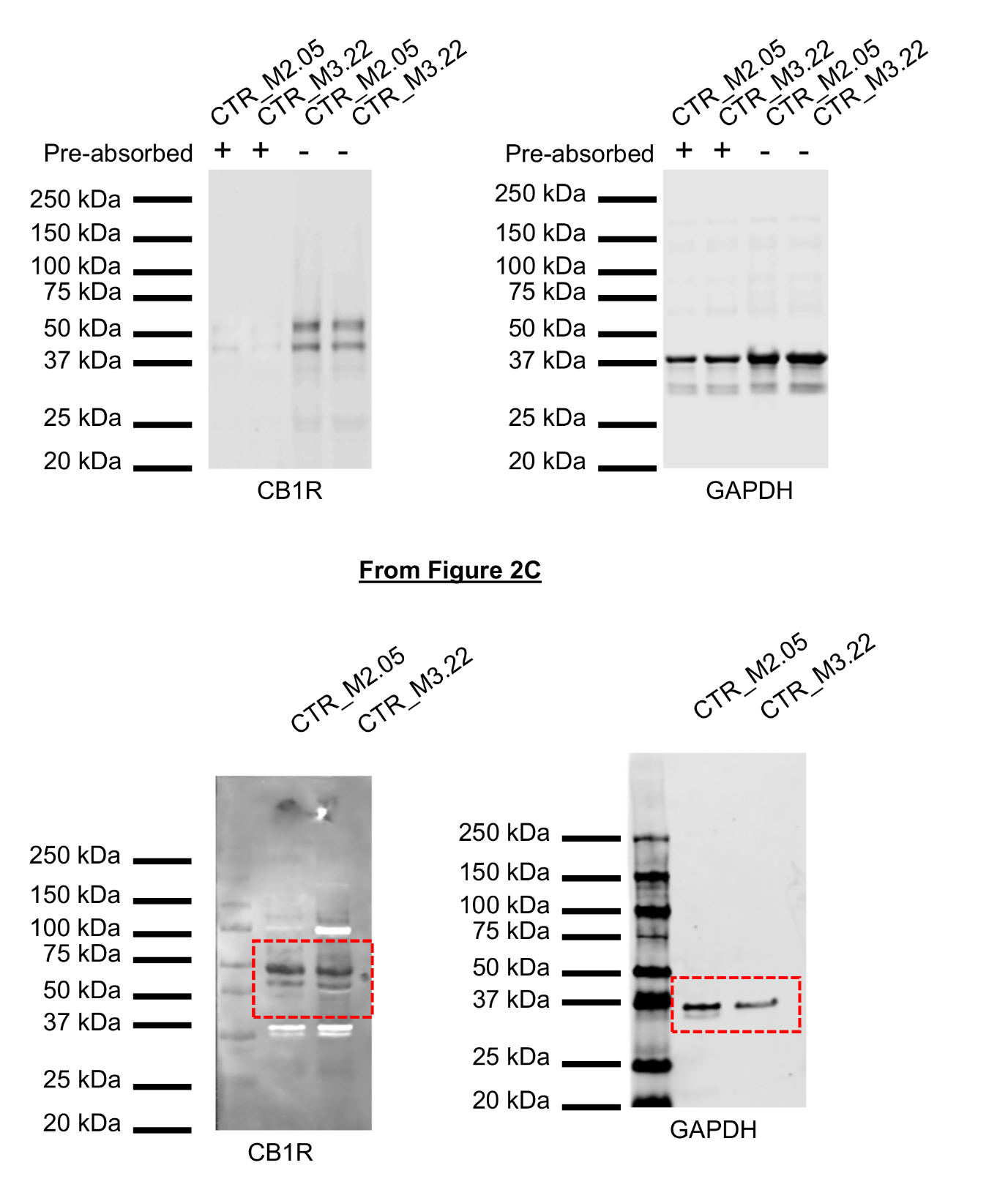

**Supplementary Figure 2.** Top**:** Pre-absorption of CB1R antibody with antigenic peptide abolishes bands around 53kDa, predicted molecular weight of CB1R protein. Bottom: Full length blot of western blot shown in main Figure 3C.

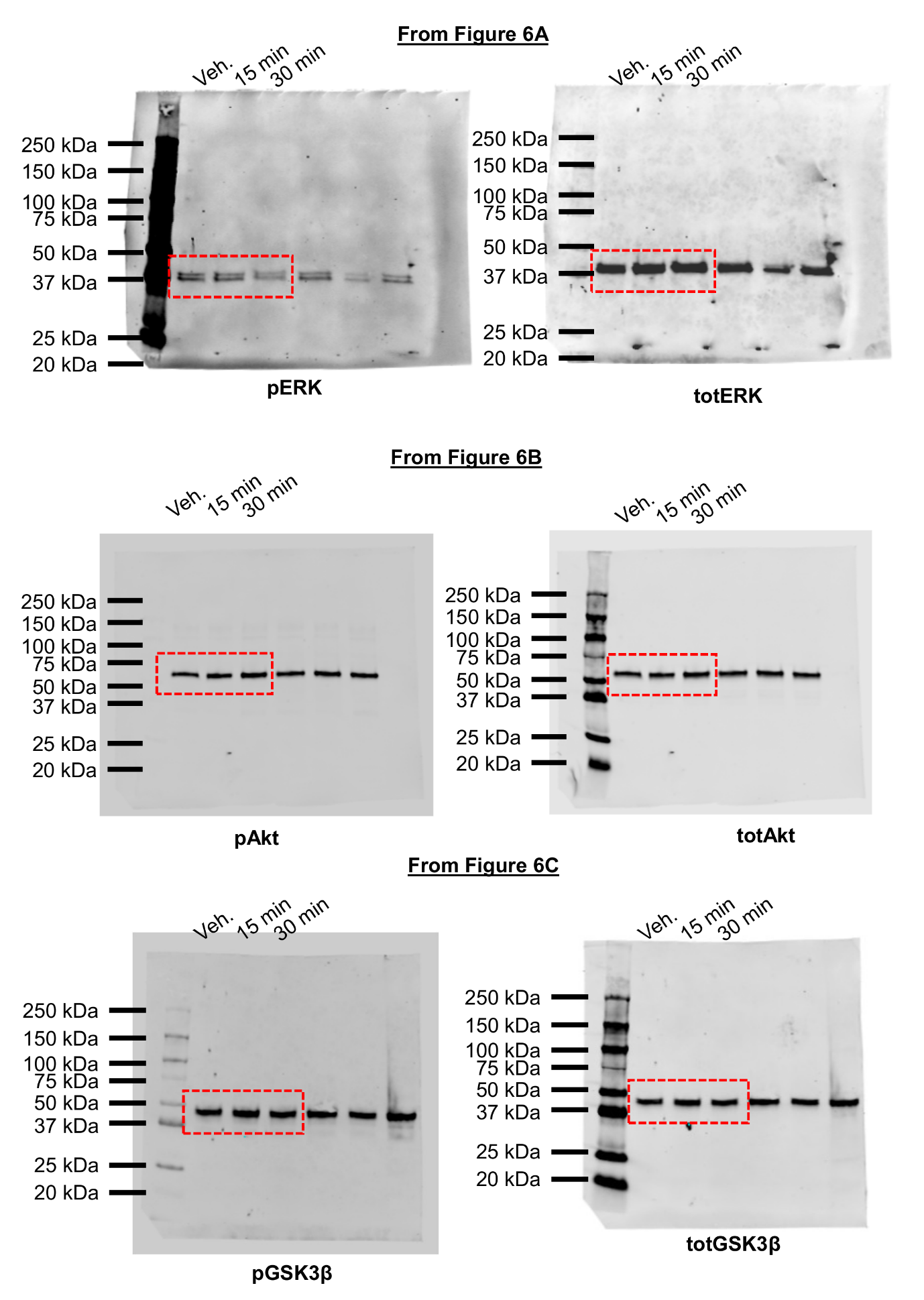

**Supplementary Figure 3.** Full length blot of western blot shown in main Figure 6 A to C.

**Supplementary Figure 4.** Full length blot of western blot shown in main Figure 6 D to F**.**

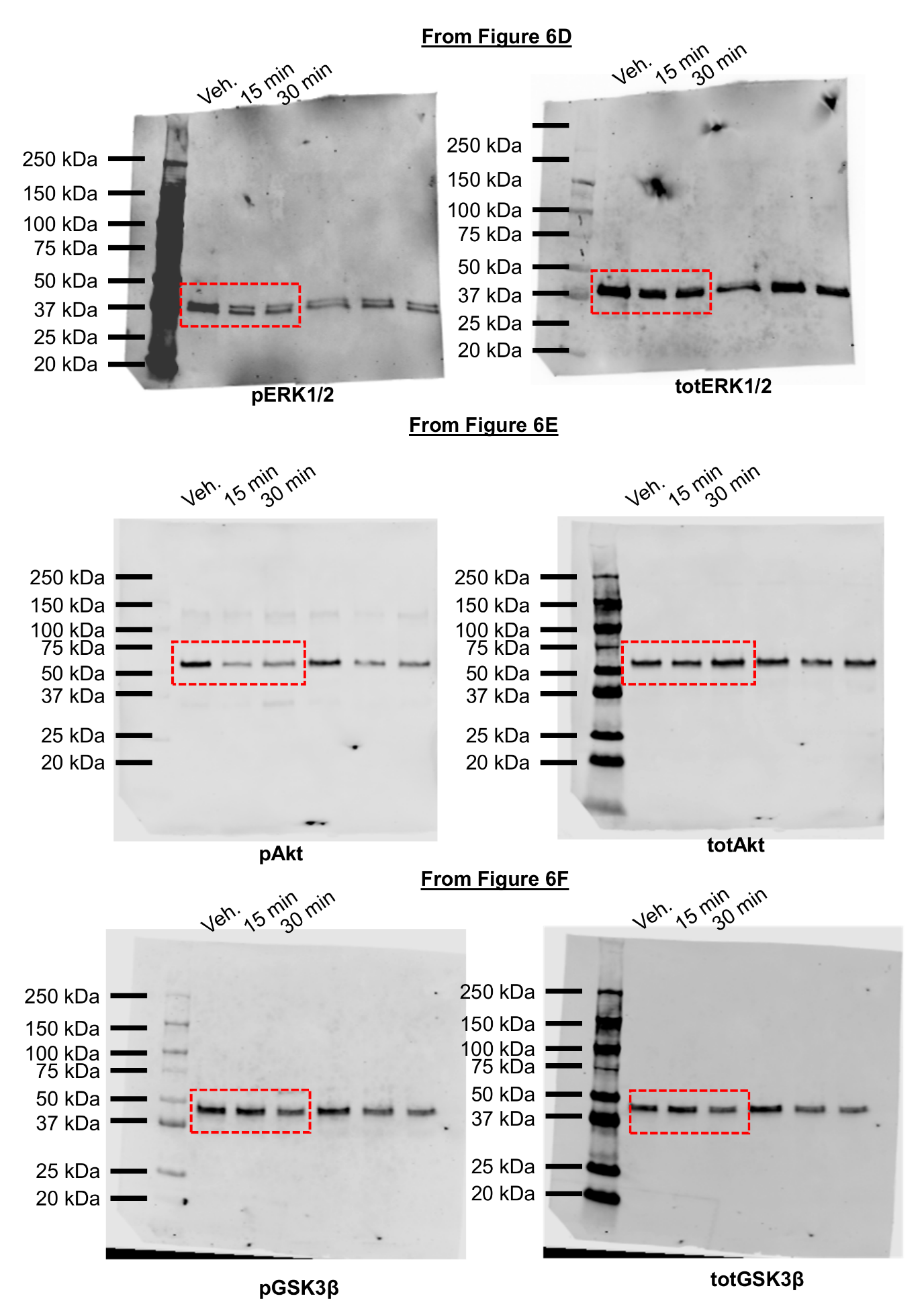

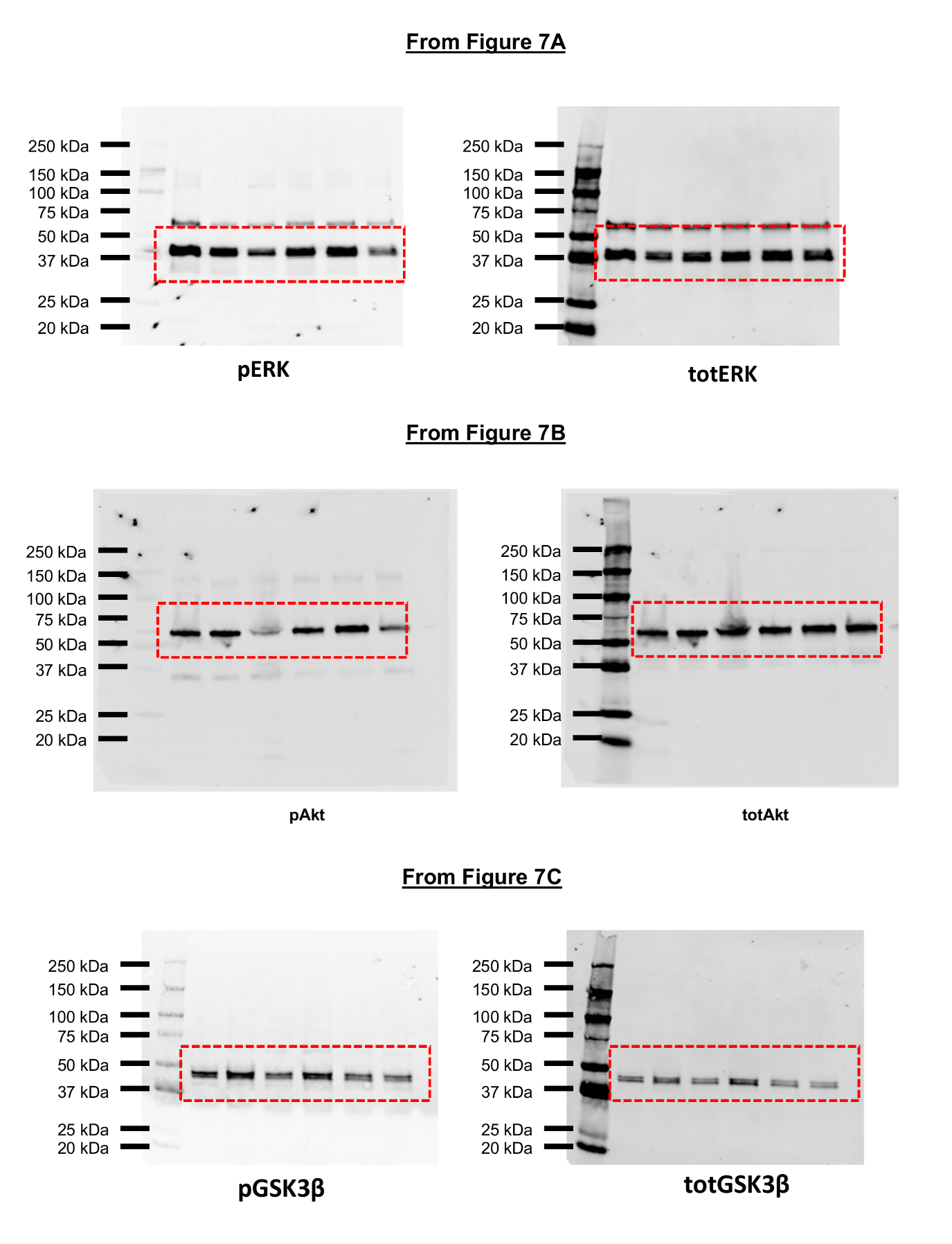

**Supplementary Figure 5.** Full length blot of western blot shown in main Figure 7A to C**.**
